# Attention and Explicit Knowledge Drive Predictive Sharpening in Early Visual Cortex

**DOI:** 10.1101/2025.10.16.682824

**Authors:** Marc Sabio-Albert, David Richter, Lluís Fuentemilla, Alexis Pérez-Bellido

**Affiliations:** Department of Cognition, Development and Education Psychology, Faculty of Psychology, University of Barcelona, Spain; Institute of Neuroscience (UBNeuro), University of Barcelona, Spain; Mind, Brain and Behavior Research Center (CIMCYC), University of Granada, Granada, Spain; Donders Institute for Brain, Cognition and Behaviour, Radboud University Nijmegen, Nijmegen, the Netherlands; Bellvitge Institute for Biomedical Research, Hospitalet de Llobregat, Spain; Department of Basic, Development and Education Psychology, Faculty of Psychology, Autonomous University of Barcelona, Spain

## Abstract

Perception is increasingly understood as an inferential process, whereby what we perceive results from the integration of sensory inputs with expectations derived from prior knowledge. Top-down predictions have been shown to alter the encoding of sensory information, from early to late stages of processing. Yet, how such predictions shape neural representations in sensory cortices remains debated. Competing accounts suggest that predictions either sharpen neural representations by enhancing selectivity or dampen activity by broadly suppressing stimulus-driven responses. In a preregistered fMRI study, we tested whether these effects depend on the level of attentional engagement and explicit knowledge of predictive associations. Using a multisensory fMRI paradigm with concurrent but independent visual and auditory probabilistic associations (75% validity) and manipulated attention, we investigated predictive effects in human early visual cortex. Consistent with prior work, expected visual stimuli elicited reduced BOLD activity. Critically, sharpening of expected visual stimuli occurred exclusively when visual inputs were attended and the concurrently presented auditory inputs expected. In addition, the magnitude of the sharpening of visual representations correlated positively with participants’ explicit knowledge of the visuo-predictive associations. These findings highlight the key roles of attention and explicit knowledge in promoting predictive sharpening and underscore the need to study predictive processing in more ecologically valid, multisensory contexts.

## Introduction

A growing body of evidence suggests that perception is fundamentally an inferential process, wherein prior expectations about the environment are merged with incoming stimuli to shape what we see and hear (Clark, 2013; de Lange et al., 2018; Girshick et al., 2011; Heilbron & Chait, 2018; Stocker & Simoncelli, 2006; Walsh et al., 2020). Consequently, expected stimuli tend to be identified faster (Chang et al., 2015; Pinto et al., 2015), with greater precision (Meijs et al., 2018; Teufel et al., 2018; Wyart et al., 2012), and typically evoke attenuated neural responses in sensory cortices compared to unexpected events (Han et al., 2019; Kaposvari et al., 2018; Meyer & Olson, 2011; Richter & de Lange, 2019), a phenomenon known as expectation suppression (ES). ES has been classically associated with a reduction in prediction error magnitudes for correctly anticipated stimuli. According to this interpretation, when predictions correctly anticipate sensory inputs, minimal mismatches occur between the observer’s internal model of the state of the environment and the input, reducing the resulting prediction errors and consequently lowering metabolic activity.

Predictions are also known to modulate the encoding of perceptual information at the representational level (de Lange et al., 2018; Press et al., 2020). Nevertheless, there is ongoing debate about how exactly predictions shape these representations. Some studies report that neural responses to expected stimuli in sensory areas are "sharpened" compared to unexpected inputs (Bell et al., 2016; Garlichs & Blank, 2024; González-García & He, 2021; Kok et al., 2012; Yon et al., 2018). The sharpening account posits that predictions modify the neural encoding of sensory inputs, acting as a noise suppression mechanism (figure 1A, left). That is, neurons tuned to the anticipated stimulus maintain—or even enhance—their gain towards the expected sensory inputs, while neurons selective for other unexpected features are suppressed. However, other studies provide evidence supporting an alternative mechanism known as "dampening” (Kumar et al., 2017; Meyer & Olson, 2011; Richter et al., 2018, 2022; Yan et al., 2023). Dampening aligns with the ‘explaining away’ idea behind predictive coding theory (Friston, 2005; Rao & Ballard, 1999) and suggests a broad suppression of neurons that preferentially respond to the predicted stimulus (figure 1A, right), resulting in an overall reduction in population responses. Typically, evidence for these two accounts has been obtained using multivariate decoding approaches: enhanced multivariate decoding for expected compared to unexpected stimuli has been interpreted as evidence for sharpening, whereas the opposite pattern as evidence for dampening.

**Figure 1.**
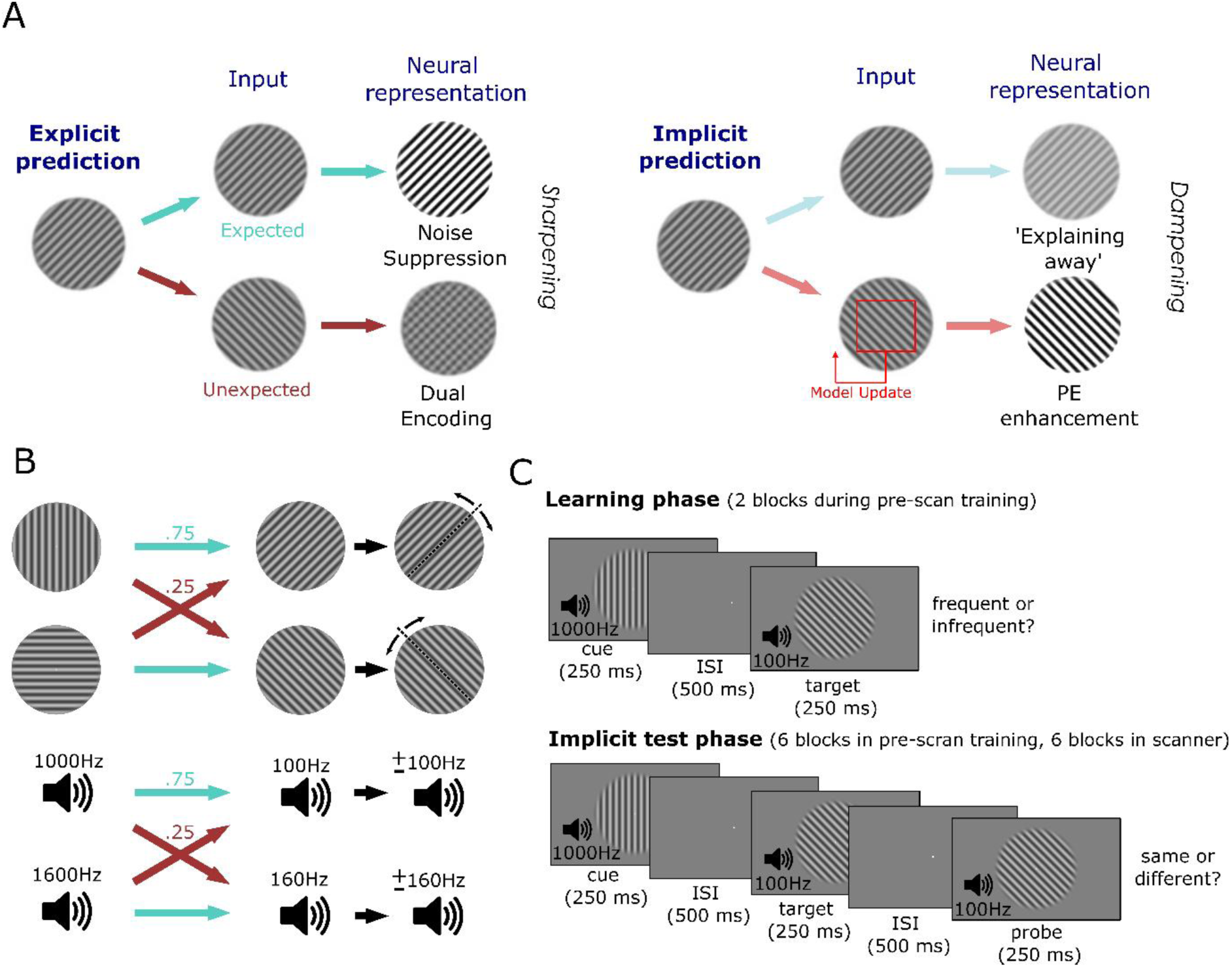
(A) Schematic overview of proposed neural mechanisms underlying predictive sharpening and dampening. Classical accounts emphasize the processing of validly predicted inputs: sharpening results from the suppression of units that are tuned away from the prediction while dampening, on the contrary, from the suppression of expected inputs. We extend these frameworks by proposing that explicit predictions yield sharpening-like patterns, as top-down preactivations are mixed with unexpected features in the sensory input. Implicit predictions, on the other hand, are resolved locally with minimal attentional needs, but prediction errors are highlighted to update internal models of the environment. (B) Experimental stimuli and probabilistic transitions. We used Gabor patches and auditory pure tones, that were presented simultaneously during the task. Within each modality, the leading stimulus predicted with 75% validity the trailing stimulus. (C) Phases of the experiment and trial sequence. The pre-scan training started with the explicit learning phase. In that phase we presented pairs of visual and auditory stimuli. In the visual blocks, participants reported whether the presented visual pairs were frequent or infrequent (ignoring auditory pairs), and in the auditory blocks the same thing but for auditory pairs. The block modality order was randomized for every participant. Learning was facilitated by feedback after every response (fixation dot turning green for correct or red for incorrect responses). Also before scanning participants underwent 6 blocks of the implicit test phase (3 visual and 3 auditory in alternating order). In this phase, the pairs retaining the same probabilistic association were presented. However, the task here was to report if the probe stimulus of the attended modality was identical or different from the target stimulus: on half of the visual attended trials the orientation of the probe Gabor slightly deviated from the standard orientations of targets (45**°** or 135**°**). Similarly, in auditory attended blocks, the probe tone had a slight deviation in its frequency in regards to the standard targets (100Hz or 160Hz). This was the main part of the experiment and the one on which we centered our neuroimaging analyses. Immediately after completing the six pre-scan training blocks, participants were led to the scanner and completed six more blocks.

In a preregistered report (https://osf.io/qf4t9), we proposed the hypothesis that the explicit versus implicit nature of the predictive associations may partially explain these divergences. The observation that motivated this hypothesis is that prediction studies reporting dampening of neural representations (Kumar et al., 2017; Meyer & Olson, 2011; Richter et al., 2018, 2022; Yan et al., 2023) tend to employ paradigms involving many stimuli combined via large transitional probability matrices, potentially hindering participants’ explicit awareness of associations. In contrast, studies reporting representational sharpening have often used paradigms with a significantly smaller set of predictive cues and targets (Bell et al., 2016; Garlichs & Blank, 2024; Kok et al., 2012; Yon et al., 2023). Such designs may facilitate the development of explicit awareness of the associations compared to paradigms that use larger transitional probability matrices. This, in turn, should favour the formation of explicit, robust predictions, particularly when the stimuli are task-relevant. However, because these studies rarely assess whether participants develop explicit knowledge of the associations, it remains unknown whether such knowledge might determine whether sensory inputs are sharpened or dampened. In a recent study from our lab (Sabio-Albert et al., 2025), we used a probabilistic cueing paradigm with a limited set of stimuli and, at the end of the experiment, assessed whether participants could explicitly discriminate the previously associated stimuli. We found that certain associated pairs were explicitly recognized. Interestingly, these explicitly recognized pairs showed a stronger predictive effect than implicitly learned pairs, demonstrating that explicit awareness modulates the impact of predictions on perception.

Our hypothesis builds on the idea that just as learning can be implicit or explicit, the human brain might also anticipate sensory inputs via implicit and explicit predictions. Specifically, if a stimulus association has been explicitly learned and is relevant to the task, it can be explicitly anticipated, and its perceptual impact may differ from that of an implicitly anticipated stimulus. Implicitly learned expectations develop gradually through repeated exposure to environmental regularities via automatic statistical learning mechanisms (Conway, 2020; Reber, 2013). Accordingly, such predictions may also be automatically instantiated with minimal attentional requirements. Large prediction errors arising from significant violations of implicit predictions may trigger a reorienting of perceptual processing to inform model updating (Press et al., 2020). This process would highlight unexpected sensory features, thereby facilitating the neural representation of unexpected stimuli, characteristic of representational dampening (figure 1A, right).

In contrast, explicit predictions, following explicit learning, may be implemented by an attention-dependent mechanism (Conway, 2020; Sabio-Albert et al., 2025) with distinct neural modulations from those of implicit prediction. We hypothesized that such explicit mechanism engages more “stubborn” predictions. These evidence-resistant predictions have been identified for low-level sensory priors such as the light from above bias (Ramachandran, 1988), or for predicting the sensory consequences of one’s own actions (Yon et al., 2018, 2019). However, it is also possible that contextual, explicitly learned associations may resist updating in the face of contradictory sensory evidence. For instance, when we hold strong certainty that a specific event will occur (e.g., a traffic light turning red after yellow), it may be adaptive to maintain that expectation, even if on one rare occasion the outcome differed (e.g., the light turned green). In line with this hypothesis, Bévalot and Meyniel (2024) demonstrated that implicit and explicit predictions rely on different computations. Whereas implicitly learned predictions adhered to Bayesian principles, they found that predictions based on explicit prior knowledge relied on heuristic computations, resulting in models that were more resistant to sensory violations. We hypothesize that this suboptimal integration of likelihood and prior knowledge may be neurally implemented as a top-down preactivation of the expected stimulus in sensory regions (Kok et al., 2014, 2017). We conceptualize this preactivation as a mechanism that is beneficial when it matches the attended input, making its processing more efficient; however, if the input is unexpected, such a stubborn prediction may resist the contrasting sensory evidence and be jointly encoded with the unexpected input (i.e., the prediction error; figure 1A, left). This concurrent encoding of the unexpected input and the prediction in sensory areas would reduce the acuity of the neural representation of the stimulus, leading to the characteristic sharpening pattern.

To test our hypotheses, we adapted a multisensory probabilistic cueing design (Sabio-Albert et al., 2025) for functional magnetic resonance imaging (fMRI). While attending to either the visual or auditory modality, participants were exposed to concurrent, predictable visual (oriented gratings) and auditory (pure tones) stimuli and performed a perceptual task that was orthogonal to the predictive regularities. Following the fMRI experiment, we used an explicit recall task to assess participants’ awareness of the visual and auditory associations. Our results showed neural sharpening in the early visual cortex that was specific to attended (task-relevant) stimuli and was further accentuated by participants’ explicit knowledge of the stimulus associations.

## Materials and Methods

### Participants

Twenty-seven students from the University of Granada, all with normal or corrected-to-normal vision and hearing, participated voluntarily in the experiment in exchange for a monetary reward. Two participants were removed due to low performance in the main task (less than 60% correct responses in a 2AFC). The final sample (n=25) consisted of 15 females and 10 males, with ages ranging from 18 to 29 (M = 21.5, SD = 3.3).

The experimental protocol was approved by the bioethics committee of the University of Barcelona in accordance with the Declaration of Helsinki. All participants received general information about the project and signed an informed consent prior to the performance of the tasks.

### Stimuli

Visual and auditory stimuli were generated using PsychoPy version 2021 (Peirce et al., 2019). The visual stimuli comprised Gabor gratings with 0.7 cycles per degree, spanning 20° of visual angle, and were presented at the center of a grey background for 250 ms. Auditory stimuli consisted of pure tones generated online and delivered through headphones at a sample rate of 44.1 kHz for 250 ms. Each trial began with the presentation of a vertically or horizontally oriented grating, which predicted—with 75% validity— whether the orientation of a subsequent grating would be rotated clockwise (CW) or counterclockwise (CCW; figure 1A). Simultaneously, a 1000 Hz or 1600 Hz pure tone was played, which in turn predicted that the upcoming tone would have a frequency of either 100 Hz or 160 Hz (figure 1B). Thus, visual and auditory leading and trailing stimuli formed predictive pairs based on their transitional probabilities. We balanced the co-occurrence of these pairs across the two modalities to avoid audiovisual integration and to ensure that visual cues could not predict auditory targets (and vice versa).

### Procedure

The experiment commenced with a training session conducted outside the scanner. At the outset, participants were explicitly informed about the probabilistic associations between the leading and trailing stimuli, as their initial task was to learn these associations. Our central hypothesis was that explicit knowledge is a necessary condition for predictive sharpening. Given that in experimental contexts with relatively simple association structures (such as ours) explicit knowledge may emerge spontaneously in only a subset of participants, we aimed to homogenize the availability of explicit predictions across the entire sample. This explicit learning phase (figure 1C) consisted of two blocks of 72 trials each. At the beginning of each block, participants were instructed to respond based on either the grating pairs (visual block) or the pure tone pairs (auditory block). A visual icon of an eye or a speaker remained at the bottom of the screen during each block as a reminder. Each trial began with the presentation of a white fixation dot for a duration randomly selected from a uniform distribution between 750 and 1500 ms. Subsequently, while the fixation dot remained on the screen, the visual and auditory leading stimuli were presented for 250 ms. After both cues disappeared, the fixation dot remained for an additional 500 ms, followed by a 250 ms presentation of the visual and auditory trailing stimuli. Then, a screen displaying two response options (“frequent” and “infrequent”) along with their corresponding keys (z or m, randomized for each block) appeared and remained until the participant responded. The fixation dot turned green following correct responses and red following incorrect responses. A third response option, “weak” (spacebar key), was available to indicate catch trials, in which one of the trailing stimuli was presented with reduced contrast (for gratings) or lower volume (for pure tones). Each block included eight catch trials (four visual and four auditory). After the explicit learning phase, participants received instructions for the implicit test phase (figure 1C), which consisted of six blocks of 72 trials each (three visual blocks and three auditory blocks, interleaved). The stimuli and pair associations remained the same as in the previous phase; however, after the trailing stimuli, an additional 500 ms inter-stimulus interval (ISI) was introduced, followed by the presentation of visual and auditory probe stimuli. These probes were either identical to the trailing stimuli of the task relevant modality or exhibited a subtle deviation in orientation or frequency, and participants were required to determine whether such a deviation was present by way of a two-alternative forced choice (2AFC).

The magnitude of this deviation was controlled for each participant and sensory modality using an adaptive staircase procedure following the “3 down 1 up” rule, targeting a deviation detection rate of 79.4%. This manipulation ensured that task difficulty was balanced across participants and sensory modality blocks. Additionally, each implicit test phase block included eight catch trials (four visual, four auditory) and provided visual feedback after every response. Immediately after completing the six blocks of implicit test phase training, participants entered the scanner, where they completed six additional runs of the same task. The task was identical, with the only difference being the jittering and duration of the inter-trial intervals, which were optimized for rapid event-related fMRI designs using OptSeq (Dale, 1999). After completing the six blocks of the main task, participants underwent two 40-trial functional localizer blocks, which employed the same task but omitted the presentation of a leading stimulus, thereby preventing predictions about the (formerly) trailing stimuli. Following these two blocks (one visual and one auditory), participants exited the scanner and completed two blocks of a brief explicit recall phase. These unimodal blocks were used to assess whether participants explicitly retained the associations learned during the initial training phase. Each of the four possible pairs within a sensory modality was presented twice (8 trials per modality), and participants indicated whether the pair was “frequent” or “infrequent.” If a participant correctly categorized more than 2 of the 4 possible transitions given a cue stimulus (e.g.: 2 vertical -> CW or 2 vertical -> CCW) as frequent or infrequent, we labelled that association as learned.

### fMRI Data Acquisition

Anatomical and functional images were acquired using a 3T Siemens Magnetom Prisma_fit scanner (Siemens, Erlangen, Germany) equipped with a 64-channel head coil. High-resolution anatomical images were obtained using a T1-weighted MPRAGE sequence (TR = 2530 ms, TE = 2.7 ms, voxel size = 1.0 × 1.0 × 1.0 mm, matrix size = 176 × 224 × 224, field of view = 176 × 224 × 224 mm). Whole-brain functional images were collected using a T2*-weighted gradient echo EPI sequence (TR = 1500 ms, TE = 40 ms, flip angle = 75°, voxel size = 2.0 × 2.0 × 2.0 mm, field of view = 210 × 210 mm, 68 interleaved slices, phase encoding direction = A>>P, bandwidth = 2090 Hz/Px) with iPAT GRAPPA acceleration (factor = 4).

### fMRI Data Preprocessing

We preprocessed the functional data using FSL 6.0.0 (Jenkinson et al., 2012). Brain extraction (BET), high-pass filtering (128 msecs), slice timing correction for interleaved acquisition, and motion correction (MCFLIRT) were performed. Head movement parameters were subsequently included as nuisance regressors in the GLMs. For the univariate analyses, functional volumes were aligned to the structural T1 image using boundary-based registration (BBR) and transformed to MNI standard space using linear transformations (12 degrees of freedom). Spatial smoothing was applied using a Gaussian kernel of 4 mm. For the multivariate analyses, spatial smoothing was omitted, and all analyses were conducted in native space by aligning functional volumes from each subject across different runs to the first run using rigid body transformations (6 degrees of freedom).

### Region of Interest Definition

We created anatomically defined regions of interest (ROIs) using the FreeSurfer 6.0.0 recon-all pipeline (Fischl, 2012). V1 and V2 labels were assigned on each subject’s native cortical surface space (Hinds et al., 2008)and subsequently transformed to volumetric space. Finally, we combined V1 and V2 masks to create a single mask of the early visual cortex (EVC). A similar procedure was followed for the A1 mask, for which we delineated Heschl’s gyrus (11133), planum temporale (11136), and superior temporal gyrus (11175) using the Destrieux anatomical atlas.

In addition, we used independent functional localizers to identify voxels that were sensitive to visual or auditory stimuli. Separately for each modality, we convolved a double gamma HRF with the onsets of each stimulus during the localizers (CW or CCW visual orientation / low or high auditory frequency), along with the first temporal derivative of each regressor and 24 movement regressors (FSL’s standard plus extended motion parameters) as nuisance regressors. We then combined parameter estimates from the two localizer runs at the second level using fixed effects, and obtained significant clusters (Z > 3.1, p < .05) of voxels that responded to either stimulus. For each participant, we ranked all voxels within significant clusters based on their Z values, and defined masks with subpopulations of voxels that were maximally responsive (50 to 350 voxels in steps of 50). When clusters of a participant were too small to create a mask, we repeated the second level analyses but decreasing the Z threshold from 3.1 to 2.3. Even after this adjustment, two participants did not reach 100 voxels in their masks. Therefore, given the reduced sample size for masks smaller than the full ROI, we report the main analyses using the entire ROIs, and include the results based on the functional masks in Supplementary Materials.

### Univariate Analyses

To model the BOLD signal in the main task functional data, we fitted a voxel-wise GLMs using FSL’s FEAT. We accounted for simultaneous visual and auditory expectations, yielding four event types: visual expected – auditory expected; visual expected – auditory unexpected; visual unexpected – auditory expected; and visual unexpected – auditory unexpected. The four regressors of interest, and two additional regressors of no interest for visual and auditory catch trials, were created by convolving the event onsets with a double-gamma HRF. Event onsets were defined at the presentation of the trailing stimuli, with event durations set to 1 second (from target stimulus presentation until the end of the probe stimulus presentation). The first temporal derivative of each regressor, along with 24 movement regressors (FSL’s standard plus extended motion parameters), were included as nuisance regressors. We defined a contrast of interest as the difference between unexpected and expected events (expectation suppression, ES). Data across the six runs were combined using FSL’s fixed effects analyses, with separate regressors for the three visual-attended runs and the three auditory runs. Finally, group-level analyses were conducted using FSL’s mixed effects analysis (FLAME 1), with Gaussian random-field cluster thresholding (cluster formation threshold Z > 2.3 and cluster significance threshold p < .05) applied to correct for multiple comparisons. The significant clusters were identified using the Harvard-Oxford atlas (FSL).

## Multivariate Analyses

### Support vector machine decoding

We estimated single-trial beta maps using the least-squares separate approach (Mumford et al., 2012). This method involves defining a separate GLM for each trial, using the onset of the current trial’s stimulus as the regressor of interest and additional regressors for the remaining trials. To better isolate neural activity associated with the presentation of visual and auditory stimuli, separate GLMs were defined for each sensory modality. For each model, regressors of no interest included the onsets of target stimuli of that modality in the remaining trials (CW vs. CCW for visual, low vs. high for auditory). These regressors and their temporal derivatives were convolved with a double-gamma HRF, and parameter estimates for each trial were extracted. For each ROI, the single-trial estimates from the main task were used to train and test a linear Support Vector Machine (SVM) as implemented in scikit-learn (Pedregosa et al., 2011). A leave-one-run-out cross-validation procedure was employed: the classifier was trained on trials from five of the six runs and tested on the trials from the remaining run, with this process repeated for each run. Notably, due to the probabilistic manipulations in our task, the number of trials across conditions was unbalanced; specifically, the most common trials in each run were those in which both the visual and auditory trailing stimuli were expected (40 trials versus 8 trials in the other conditions). To mitigate biases resulting from this imbalance, we performed 200 permutations of a random stratification approach, in which 8 trials from the more numerous condition were pseudo-randomly selected for each training fold. We also ensured that stimulus identities were balanced across both modalities within the selected trials. This procedure yielded 32 trials for each training run (32 × 5 = 160 training trials). The trained classifiers were then tested on all 64 trials of the remaining test run. We applied this decoding approach to our pre-defined ROIs (EVC and A1).

### Modelling SVM predictions

To test our hypothesis regarding the sharpening and dampening of sensory representations, we departed substantially from our preregistered analysis plan. This deviation was primarily prompted by the near-chance decoding accuracy of visual information from localizer data in most participants. To address these limitations, we developed a new analysis strategy designed to more effectively test our core hypotheses. We conducted statistical analyses on two different metrics yielded by the classifiers trained and tested on the main experiment data. The first measure was decoding accuracy, which refers to the proportion of correctly predicted labels. While accuracy provides a straightforward measure of classifier performance, it is inherently binary and does not capture the degree of classification certainty. To address this, we obtained a second metric: probability values reflecting classification certainty for a specific stimulus label. For this, we applied Platt scaling (Platt, 1999), a widely used post-hoc calibration method. This approach fits a logistic sigmoid function to the raw classifier scores following an additional calibration step, in which a subset of the training data is used to estimate the transformation parameters. The result is a continuous probability score (ranging from 0 to 1) that can be interpreted as the classifier’s certainty in its prediction.

We first examined whether decoding accuracy varied across expectancy conditions (expected vs. unexpected) to test for evidence of prediction-induced dampening (reduced decodability for expected stimuli) or sharpening (enhanced decodability for expected stimuli). These effects were assessed using repeated-measures ANOVAs. To control for biases due to the imbalance between expected and unexpected trials, we applied random down-sampling: for each of the 200 permutations produced by our cross-validation procedure (see Support Vector Machine Cross-Validation), we randomly selected 8 trials from the overrepresented "visual expected – auditory expected" condition, matching the logic of our previous approach. To further explore the underlying mechanisms, we analysed the classification probabilities, which offer a graded index of representational fidelity beyond binary outcomes. Using repeated-measures ANOVAs, we assessed how classification certainty toward the presented sensory input (e.g., presented CW vs. presented CCW) was modulated by the predicted orientation (e.g., expected CW vs. expected CCW) cued in advance.

We also investigated whether these effects were influenced by explicit knowledge of the predictive associations. For this, we employed a mixed-effects linear regression model in which the logit-transformed classification probabilities were regressed on the interaction between the visual expectation (expected CW vs. expected CCW) and a binary variable indicating whether the prediction was explicitly learned. Trials were categorized as "learned" or "not learned" based on each participant’s post-scan recall performance (see Procedure). Subject-level intercepts were included as random effects to account for inter-individual variability.

Importantly, to avoid inflating the number of observations, we aggregated the classifier probabilities across the 200 permutations for each trial, computing the mean predicted probability per trial and per subject. This step was necessary because each permutation used a different subsample of training data, which led to variability in classifier predictions for the same trial and violated the assumption of independence.

Unlike the decoding accuracy analyses, this modelling approach did not require down-sampling. This is because the predictors in the regression model (expected stimulus identity and its explicit recall) were orthogonal to the overrepresentation of the “expected” condition. However, this strategy was limited to analyses that excluded the presented stimulus identity as a regressor, since including both expected and presented stimulus identities would reintroduce the same imbalance.

## Results

### Predictions did not modulate behavioural performance

We tested if participants performed the task correctly, and whether their performance was modulated by predictive information. Accuracy in visual attended runs, measured as the proportion of correct responses (group mean = .837; SD = .049), was slightly higher than in auditory attended runs (group mean = .817; SD = .05), but both measures were very close to the one targeted with the staircase procedure (.794). Visual expectations had no significant effect on accuracy (*t*(24) = 1.06, *p* = .302, *d* = .15) or reaction times (*t*(24) = -0.22, *p* = .83, *d* = .01). Similarly, auditory expectations did not significantly affect accuracy (*t*(24) = 0.42, *p* = .681, *d* = .07) or reaction times (t(24) = 1.00, *p* = .327, *d* = .04). Subsequent 2-way repeated-measures ANOVAs revealed that these effects were not contingent on the task-relevant (and thus attended) modality. The absence of behavioural facilitation by valid prediction likely arose from the delay between target stimulus and probe onset, thereby reducing the impact of the prediction at the decisional stage. Moreover, similar experimental paradigms have reported no behavioural modulations due to expectation, while demonstrating significant neural modulations (Aitken & Kok, 2022; Kok & Turk-Browne, 2018).

### Expectation suppression in visual cortex

Since we did not observe any significant behavioural effect, we tested for the existence of differences in neural activity between expected and unexpected trials at the whole brain level. We identified a significant cluster of visual ES in occipital regions (MNI: x 18, y = -100, z = 0; peak Z = 3.39, cluster level p = .001) when visual and auditory attended runs were combined (figure 2A), revealing that our experiment induced predictive processing in EVC. For auditory expectation, the pattern differed depending on the attended modality. When participants attended the visual stimuli, unexpected tones resulted in an increased activation of right frontal clusters, specifically in the middle (MNI: x = 28, y = 48, z = 2; peak Z = 3.29, cluster level p = .009) and superior frontal gyri (MNI: x = 18, y = 24, z = 48; peak Z = 3.33, cluster level p < .001) extending into the dorsolateral PFC. (figure 2B). Conversely, unexpected attended sounds attenuated BOLD responses in the medial superior frontal gyrus (MNI: x = 8, y = 54, z = 28; peak Z = 3.68, cluster level p < .001) and in the left angular gyrus (MNI: x = -52, y = -54, z = 20; peak Z = 3.62, cluster level p = .007) (Fig 2C). These results provide initial evidence that participants effectively learned the sensory associations. Visual expectations modulated early visual processing, as indexed by ES in the EVC. We also expected to find auditory ES in A1, but the pattern of brain activity we found instead, involving the frontoparietal network, may be related to attentional dynamics. For instance, while participants were attending to the visual stimuli, the recruitment of the observed right lateralized frontal areas in response to unexpected sounds may indicate a transient reorientation of attention (Kim, 2013).

**Figure 2.**
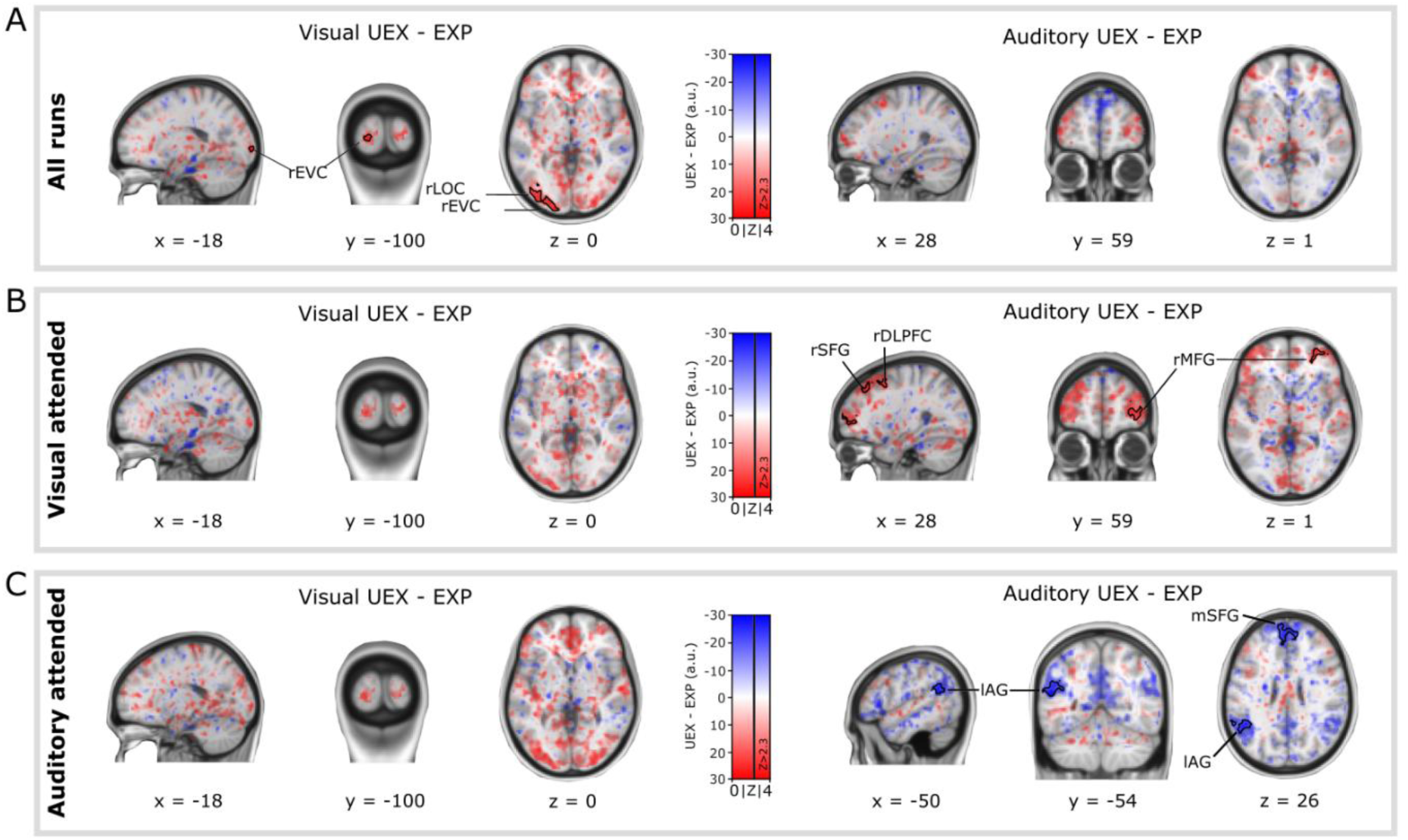
(A) Whole-brain contrast of visual (left) and auditory (right) expected vs. unexpected trials in all runs. We observed significant visual ES in occipital regions (MNI: x = 18, y = –100, z = 0; 358 voxels; peak Z = 3.39; cluster-level p = .001) (B) Whole-brain contrast of visual (left) and auditory (right) expected vs. unexpected trials in visual attended runs. Unexpected auditory stimuli elicited increased activation in right frontal regions, including the middle frontal gyrus (MNI: x = 28, y = 48, z = 2; 254 voxels; peak Z = 3.29; p = .009) and superior frontal gyrus extending into the dorsolateral prefrontal cortex (MNI: x = 18, y = 24, z = 48; 411 voxels; peak Z = 3.33; p < .001). (C) Whole-brain contrast of visual (left) and auditory (right) expected vs. unexpected trials in auditory attended runs. Unexpected sounds led to attenuated BOLD responses in the medial superior frontal gyrus (MNI: x = 8, y = 54, z = 28; 1195 voxels; peak Z = 3.68; p < .001) and the left angular gyrus (MNI: x = –52, y = –54, z = 20; 251 voxels; peak Z = 3.62; p = .007).

### No general modulation of neural representations by expectation

Next, we set out to test our central hypothesis that predictions influence sensory patterns differently depending on whether the stimuli are attended or unattended. We applied SVMs on single-trial activity patterns from EVC and A1 ROIs to classify presented visual and auditory stimulus identity, respectively. We then assessed classification performance as a function of expectation and attention towards the stimuli.

We obtained above-chance decoding in both A1 and EVC across all functionally defined mask sizes (figure S1, table S1). As expected, attention towards the visual stimuli enhanced decoding in EVC for most voxel mask sizes (figure S1, table S2). For the next decoding analyses, we focused on the whole ROI since, when selecting masks based on the number of active voxels in the functional localizer, two participants had to be excluded (see functional ROI definition). Results for other EVC mask sizes are shown in figure S2 and tables S3, S4.

To test the hypothesis that predictive sharpening and dampening take place for attended and unattended stimuli, respectively, we performed 2-way repeated-measures ANOVAs separately on the decoding accuracy using whole ROI masks for each modality. The ANOVA for visual decoding, with factors of visual attention (attended vs. unattended) and visual expectation (expected vs. unexpected) revealed that visual attention did not significantly modulate decoding accuracy (figure 3A; mean attended: 0.57 vs. mean unattended: 0.55; main effect: *F*(1,24) = 3.73, *p* = .066, partial *η^2^* < .001), and neither did visual expectation (main effect: *F*(1,24) < 0.001, *p* = .99, partial *η^2^* < .001). Finally, there was no indication of an interaction between attention and expectation (*F*(1,24) = 2.56, *p* = .123, partial *η^2^* < .001). We also run one-tailed repeated measures t-tests to directly test our directional hypotheses. We did not observe either sharpening effects of visual inputs in visual attended runs (*t*(24) = -1.25, *p* = .111, one-tailed, *d* = -.16) or dampening in auditory attended runs (*t*(24) = .84, *p* = .205, one-tailed, *d* = .17). Therefore, this initial set of analyses did not provide evidence in favour of our hypothesis that sharpening and dampening depend on attention.

**Figure 3.**
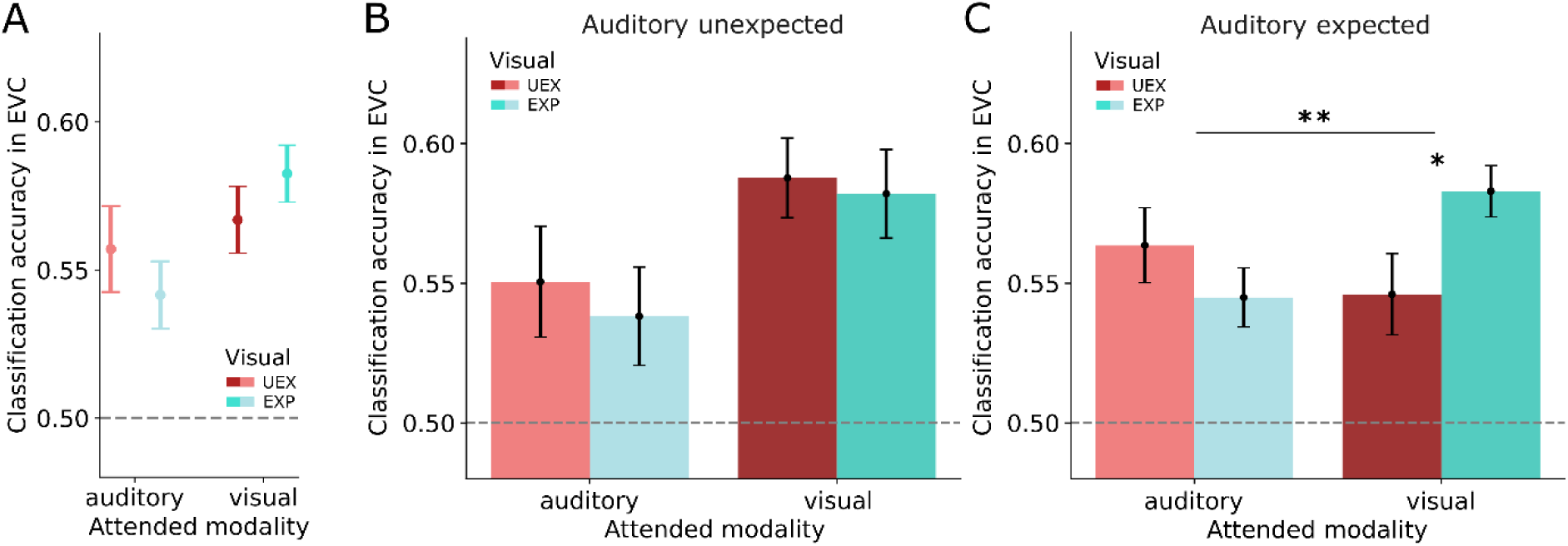
(A) Classification accuracy in EVC based on the attended sensory modality, and visual expectation. (B) Classification accuracy in EVC based on attention and visual expectation, in trials where the auditory stimulus was unexpected. (C) Classification accuracy in EVC based on attention and visual expectation, in trials where the auditory stimulus was expected. * *p_FDR_* < .05 ** *p* < .01

The same was the case in the auditory modality: attention towards auditory stimuli improved decoding accuracy (figure S3; mean attended: .61 vs mean unattended: .59; main effect: *F*(1,24) = 4.88, *p* = .037, partial *η^2^* = .02). However, auditory expectation had no reliable effect (main effect: *F*(1,24) = .3, *p* = .589, partial *η^2^* < .001) and there was no interaction between attention and expectation (*F*(1,24) = .07, *p* = .795, partial *η^2^* < .001). All subsequent analyses refer to the visual modality, but the same results for the auditory modality can be found in figure S3 and tables S5, S6.

### Sharpening of visual representations in EVC is contingent on attention and auditory expectation

In the initial analyses above, attention did not significantly impact the encoding of expected or unexpected sensory inputs. However, this null effect might be explained by how concurrent auditory expectations are processed. Specifically, in a previous study (Sabio-Albert et al., 2025) using a similar experimental design, we found that unexpected stimuli in the unattended sensory modality disrupted predictive processing in the task-relevant modality. We interpreted this effect as an interference on predictive processing triggered by expectation violations outside the focus of attention. This is further supported by the increased frontal activation of frontal regions which we identified in auditory unexpected trials during the visual runs in the current task (figure 2B). Therefore, we investigated whether visual predictive processing was contingent on concurrent auditory expectations by conducting two-way repeated measure ANOVAs on classification accuracy in EVC for auditory unexpected and expected trials separately, with attended modality and visual expectation as factors.

When auditory stimuli were unexpected, the effects of visual expectation did not change as a function of attention (figure 3B; interaction: *F*(1,24) = .05, *p* = .83, partial *η^2^* <.001) We conducted post-hoc contrasts, which did not reveal any dampening effect in the absence of attention (*t*(1,24) = .46, *p_FDR_* = .326, one-tailed, *d* = 0.11), nor sharpening in visual attended runs (*t*(1,24) = .3, *p*_FDR_ = 0.62, one-tailed, *d* = .05). However, when auditory stimuli were expected there was a significant interaction between attended modality and visual expectation (figure 3C; *F*(1,24) = 11.49, *p* = .002, partial *η^2^* = .026). Post-hoc tests revealed that the interaction was driven by a sharpening-like effect in visual attended runs (*t*(1,24) = -2.25, *p*_FDR_ = .033, one-tailed, *d* = -0.4) and no significant modulation of neural representations by expectation in unattended runs (*t*(1,24) = -1.16, *p*_FDR_ = .26, one-tailed, *d* = 0.23). These results suggest that representational sharpening may be contingent on attending the predicted visual input, but that this process can be disrupted by concurrent surprising events that transiently capture attention.

### Prediction-dependent biases in visual representations

Next, we turned to the probabilistic output of our SVMs, which provides a continuous measure of classification certainty between the two possible ‘decisions’ (CW and CCW orientations). This continuous metric (in contrast to classification accuracy obtained from the binary outcome of the classifications) allowed us to assess separate contributions of the actual sensory inputs and the expected sensory inputs on the classifier performance. That is, when decoding visual orientations in EVC the classifier should reflect, by definition, stronger evidence in favour of the visual stimulus (e.g. greater ‘CW’ evidence in trials in which CW gratings were presented). On the other hand, according to our hypothesis that the brain is also encoding a representational template of the expected orientation, this evidence should be biased by the visual expectation. In other words, regardless of the stimulus input, the probabilistic classifier output should also, on average, predict the expected stimulus.

To test this, we performed a three-way ANOVA on the probability assigned to classifying CW orientations (with values below 0.5 indicating stronger evidence for CCW; see Modelling SVM predictions) as a function of visual stimulus (presented CW vs. CCW), visual expectation (CW vs. CCW) and auditory expectation (expected vs. unexpected) as factors. As anticipated, visual stimuli presentations greatly influenced the classification probability metric (main effect: *F*(1,24) = 14.97 *p* < .001, partial *η^2^* = .384), and crucially, there was a significant interaction between visual expectation and auditory expectation (figure 4A; *F*(1,24) = 5.43, *p* = .028, partial *η^2^* = .184). This interaction suggests that the effects of visual expectations were contingent on whether the auditory stimuli were expected, reinforcing our previous finding on classification accuracies. In order to unpack this interaction, we ran main effect analyses. This revealed that when auditory stimuli were unexpected, visual expectations did not bias the classifier predictions (F(1,24) = 0.25, p = .62, partial *η^2^* < .001). In contrast, when auditory stimuli were expected, visual expectations significantly modulated responses (F(1,24) = 8.7, p = .007, partial *η^2^*= .039). That is, CW and CCW inputs were more likely to be classified as CW when the expected stimulus was CW compared to when the expected stimulus was unexpected and vice versa. These results suggest that visual representations in EVC not only reflected the presented stimulus but were also biased toward the predicted orientation. In other words, neural representations partially resembled the expected stimulus feature even when this stimulus was not physically present. This prediction-induced bias emerged only when auditory expectations were confirmed, suggesting that predictive templates in the visual cortex may shape sensory representations only when the broader cross-modal predictive context is coherent.

**Figure 4.**
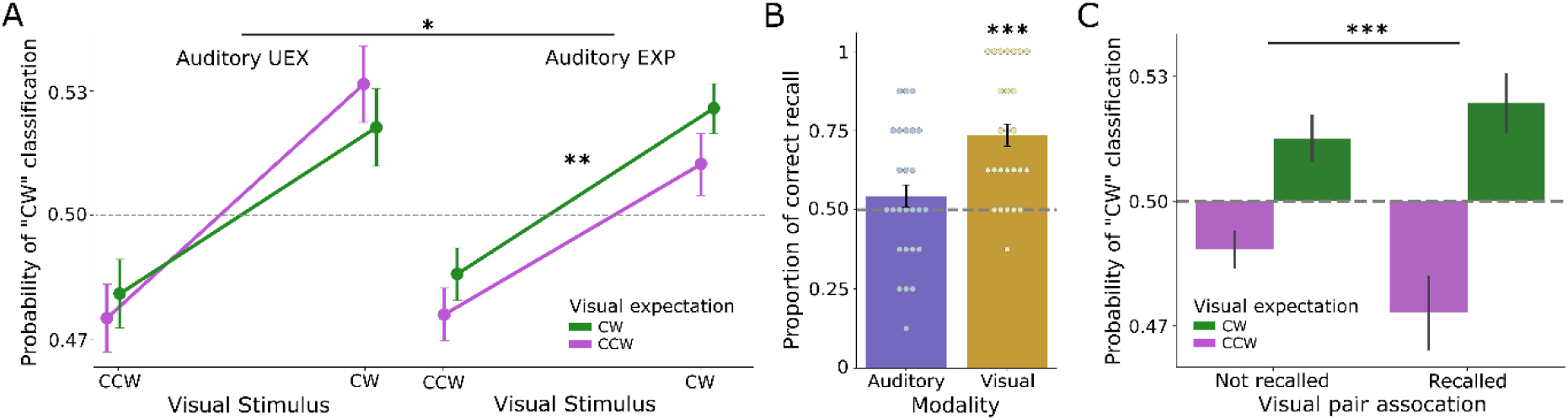
(A) Probability of “CW” classification of a linear SVM from EVC activity, based on the predicted and the presented orientations. (B) Proportion of correct recall in the final explicit recall phase, in which participants reported the auditory and visual predictive associations. The dots correspond to the accuracy of each participant for all 8 responses in each sensory modality(C) Probability of “CW” classification during visual runs based on the predicted orientation and whether the participant demonstrated explicit knowledge about the predictive association in the explicit recall phase (only auditory expected trials).

### Explicit knowledge amplifies the sharpening of visual representations

Finally, we assessed participants’ explicit knowledge of the predictive associations and its effect on neural representations. A binomial test indicated that participants on average correctly recognized the visual associations above chance (figure 4B; mean accuracy: 73.6%; *p* < .001). In contrast, recognition of auditory associations did not significantly differ from chance (mean accuracy = 54.3%, *p* = .238). This lack of significant recognition for the auditory pairs may be related to the absence of significant sharpening effects in A1, although we refrain from attributing it to that factor alone.

Following our preregistered hypothesis, we reasoned that a better long-term explicit awareness of the associations should result in greater predictive sharpening. To test this, we fit a mixed-effects linear regression to all auditory expected trials in visually attended runs, modelling the probability of CW classification (see Modelling SVM predictions) as a function of visual expectation (CW vs. CCW) and explicit knowledge (recalled vs. not recalled). A significant interaction between explicit knowledge and visual expectation (Z = 5.07, p < .001; 95% CI [.023, .053]) indicated that the bias toward the expected orientation was larger when participants explicitly anticipated? which orientation was most likely (Fig 4C). These results show that explicit knowledge of the learned associations strengthens prediction-induced biases in EVC, suggesting that consciously maintained expectations may enhance the top-down templates shaping sensory representations.

## Discussion

Perception is often understood as a form of Bayesian inference, where prior expectations combine with sensory input to shape neural responses. Yet predictions can alter representational format in two opposing ways: “sharpening,” which increases the discriminability of expected features, and “dampening,” which is thought to broadly attenuate responses to predicted inputs. The goal of this study was to test our preregistered hypothesis (https://osf.io/qf4t9) that predictive dampening of sensory representation results from implicit predictions of unattended stimuli, and that sharpening is driven by explicit knowledge of attended sensory predictions. To adjudicate between these accounts, we used a multisensory probabilistic cueing paradigm while recording fMRI. Participants attended either vision or audition while viewing oriented gratings and hearing pure tones with explicitly learned within-modality contingencies, allowing for the prediction of upcoming sensory inputs. We then quantified representational outcomes of prediction using multivariate decoding.

Our results provide consistent evidence that predictive sharpening–like effects in early visual cortex (EVC) were gated by attention and enhanced by explicit knowledge, as reflected in stronger multivariate decoding of fMRI activity for expected compared to unexpected visual stimuli in early visual areas. This occurred only when participants attended to the visual inputs and when the auditory information was predictable. While previous studies have reported similar predictive sharpening (Bell et al., 2016; Garlichs & Blank, 2024; González-García & He, 2021; Kok et al., 2012; Yon et al., 2018), our results extend this literature by showing that the effect was further amplified when participants possessed explicit knowledge of the predictive associations.

In this work, we conceptualize sharpening as the consequence of a representational interference between the encoding of sensory inputs and predicted inputs in EVC during unexpected trials. At the neural level, learned predictive associations have been shown to bias the content of neural activity long after predictions cease to be reliable (Yon et al., 2023), when expected inputs are omitted (Aitken et al., 2020), or by inducing pre-stimulus templates of expected features (Kok et al., 2014, 2017). Such representational biases may lead to competing sensory representations between predicted and presented (mismatching) inputs, resulting in mixed representations that are more difficult to identify as a specific stimulus (Kok et al., 2017).

Our finding that explicit knowledge increased predictive biases in the classification of visual inputs aligns with our hypothesis that explicit predictions might be more resistant, to sensory evidence, therefore inducing greater mismatch when visual inputs are unexpected. Such explicitly evoked predictions could be implemented through a top-down controlled attentional mechanism. Attention has been demonstrated to exert effects consistent with sharpening, such as increased perceptual sensitivity (Carrasco et al., 2000; Wyart et al., 2012) and enhanced gain in neural units selectively tuned to relevant features (Corbetta et al., 1990; Martinez-Trujillo & Treue, 2004; Maunsell, 2015; Treue, 2003). For that reason, we hypothesized that attention is necessary for predictive representational sharpening.

Conversely, other accounts interpret predictive sharpening as top-down noise suppression, whereby sensory inputs that do not match predictions are cancelled (de Lange et al., 2018). Although sharpening as noise suppression and sharpening via simultaneous encoding of mismatching representations could, in principle, co-exist (Fig. 1A; taking place for expected and unexpected inputs, respectively), our pattern of results appears to be more consistent with the latter. Specifically, the observed difference in decoding accuracy between expected and unexpected orientations (Fig. 3C) seemed to be driven by decreased accuracy for unexpected stimuli, rather than increased accuracy for expected stimuli, relative to the decoding accuracy in other attended visual conditions. Nonetheless, this remains an observation, as the present task was not designed to formally disambiguate between these mechanisms.

Although we did not plan to test how auditory predictions shape the encoding of sounds, we performed complementary exploratory analyses (Fig. S3 and tables 5-6). We found no evidence of auditory ES in the auditory cortex; however, several clusters related to predictive processing emerged in higher-level regions, indicating that participants were sensitive to auditory prediction violations. When participants attended to the auditory modality, we observed greater activation of DMN regions for expected sounds compared to unexpected ones. Given that this network has been classically viewed as anti-correlated with task engagement (Fox et al., 2005), this finding suggests that during expected trials, participants simply performed their task and waited for the second tone to make a comparison. In contrast, auditory prediction violations in unexpected trials may have elicited a surprise response accompanied by increased alertness (i.e., a reduction in DMN activity). Conversely, when attention was focused on the visual modality, we found right lateralised frontal activation elicited by unexpected tones compared to expected ones, including brain regions that are part of the ventral attention network (Corbetta & Shulman, 2002; Kim, 2013), known to activate in response to salient stimuli. This result is consistent with prior findings of auditory predictive processing during purely visual tasks (Parmentier et al., 2022; Vasilev et al., 2023). These observations lead us to consider that unexpected sounds may disrupt the top-down modulatory processes hypothesized to underlie perceptual sharpening by causing a withdrawal of attention from the visual stimuli. Behavioural data from a previous study by our group (Sabio-Albert et al., 2025) is consistent with this view, showing that target expectations ceased influencing behavioural performance when unattended stimuli were unexpected. Thus, we speculate that when unattended auditory stimuli are correctly predicted, visual predictions are implemented as if no competing stimuli were present. However, when an auditory stimulus is unexpected, it may transiently capture attentional resources—resources that might otherwise support explicit visual predictions—thereby disrupting visual prediction. This interpretation aligns with our results showing that only attended stimuli resulted in representational sharpening. It should be noted that the overall decoding accuracy for visual stimuli in EVC did not significantly drop relative to the auditory expected conditions. This indicates that the sudden unexpected sound may have impeded the expectation-dependent sharpening of visual representations by diverting attention, but because this diversion was brief, the encoding of visual stimuli remained largely intact. Overall, our results suggest a complex interplay between predictions across sensory modalities, warranting further research, particularly in more ecologically valid settings.

Contrary to one of our preregistered predictions, we did not observe dampening effects for unattended stimuli. One possible explanation is that unexpected events in our experiment did not elicit enough surprise to trigger the model-updating mechanisms proposed by Press et al. (2020). The limited number of possible stimuli and their simplicity (i.e.: Gabor patches and pure tones) distinguishes our experiment from others which did find evidence of expectation-induced dampening (Kumar et al., 2017; Meyer & Olson, 2011; Richter et al., 2018, 2022; Yan et al., 2023), wherein larger sets of more complex stimuli lead to a more dynamic, less predictable sensory environment. Thus, it is possible that in our experiment, the unexpected stimuli could always be “expected” to a certain degree, as there were only two possible outcomes with a 75% and 25% chance of appearance.

In our results, we observed visual sharpening, albeit to a lesser extent, when participants lacked significant explicit awareness of the predictions. This might be attributed to a limitation of our study given that our measure of explicit knowledge was a binarization (learned vs. not learned) with limited sensitivity, as it was based on a small number of decisions. Therefore, it may have failed to capture subtler gradations in the strength of the learned associations. One possibility is that explicit knowledge should be more accurately conceptualized as a continuum, ranging from implicit associations to progressively stronger, declarable ones (Batterink et al., 2015; Bertels et al., 2012; Cleeremans, 2011). Thus, sub-threshold associations may still fulfil similar roles as fully declarable ones, though less effectively. Future studies could mitigate these limitations by including measures such as confidence reports.

## Conclusion

In summary, sharpening of visual representations for expected stimuli was only observed when these inputs were attended and simultaneous, task-irrelevant auditory stimuli were expected. Combined, our results suggest that predictive processes in our inherently multisensory world involve complex interactions between prior knowledge across sensory modalities, underscoring the need for integrative accounts of predictive processing that span modalities. Furthermore, assessing classification probabilities (rather than accuracy) we showed that sharpening was amplified when participants possessed explicit knowledge of the predictive associations. Together, these findings highlight that predictive processes are a context-sensitive interplay between attention, cross-modal expectations, and explicit knowledge that jointly determine how sensory information is represented.

## Acknowledgments

This work was supported by the Spanish Ministerio de Ciencia, Innovación y Universidades, which is part of Agencia Estatal de Investigación (AEI), through the projects RTI2018-100977-J-I00 and RYC2022-037652-I to A.P.B. and the projects PID2019-111199GB-I00 and PID2022-140426NB-I00 to L.F. (funded by MCIN/AEI/10.130 39/501100011033/ and FEDER a way to make Europe). We thank CERCA Programme/Generalitat de Catalunya for institutional support.

