## Supplementary figures for "Attention and Explicit Knowledge Drive Predictive Sharpening in Early Visual Cortex"

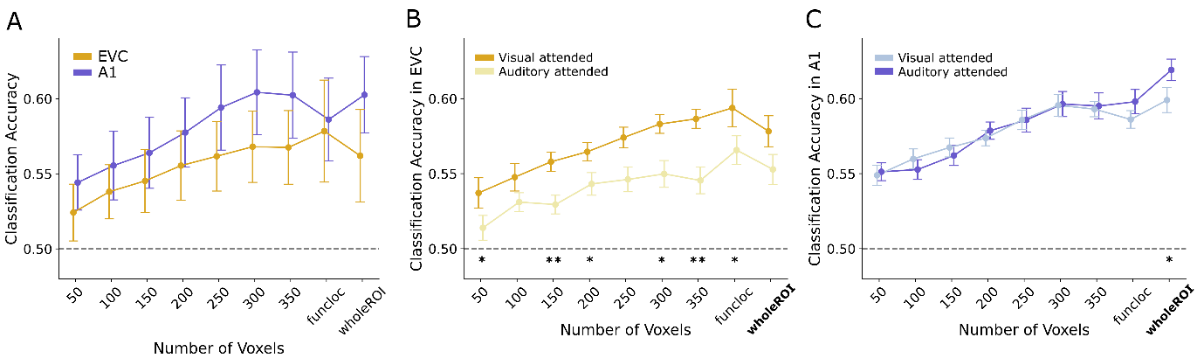


**Figure S1 (A)** Classification accuracy in EVC (yellow) and A1 (purple) for different ROI mask sizes. The accuracies were better for the auditory decoding in A1, but they were above chance in all cases. (B) Classification accuracy of presented orientation based on attended modality, using different EVC voxel mask sizes. * $p < 0.05$ ** $p < 0.01$ (C) Classification accuracy of presented auditory frequency based on attended modality, using different A1 voxel mask sizes. * $p < 0.05$

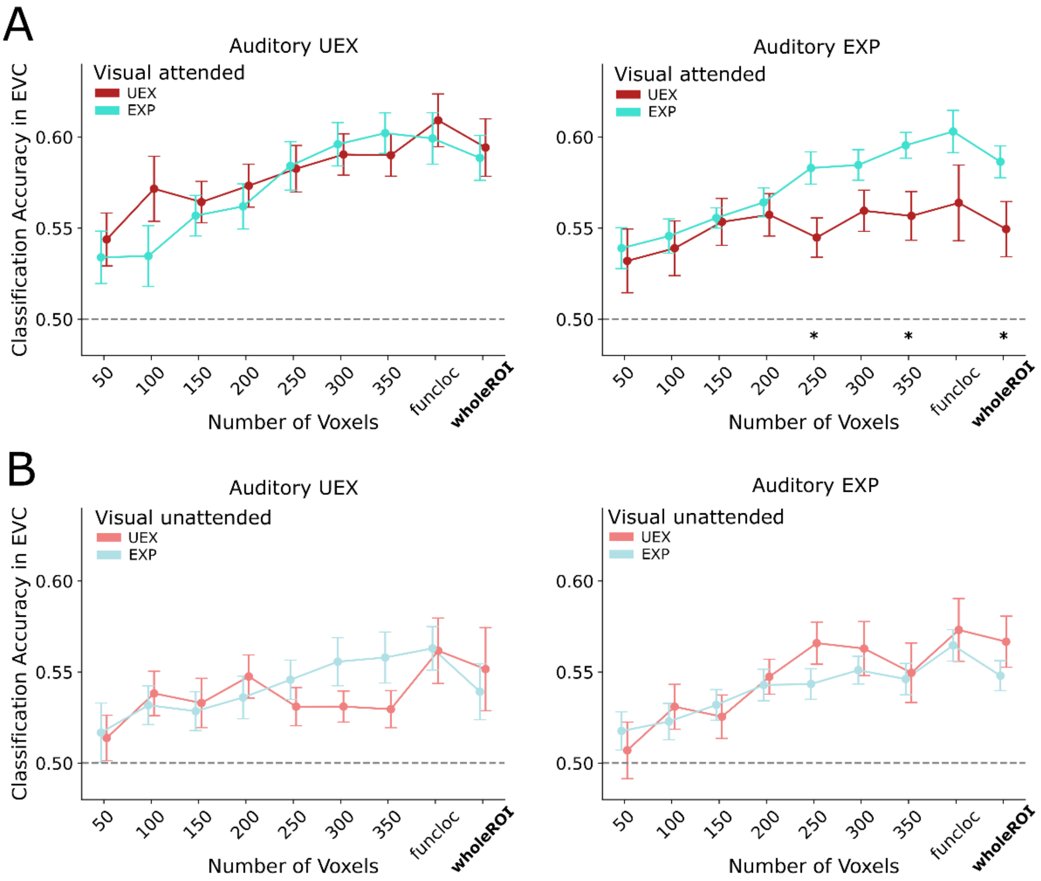

**Figure S2** (A) Classification accuracy using different EVC mask sizes in visual attended runs, based on visual expectation. The left plot shows auditory unexpected trials, and the right plot auditory expected trials. (B) Classification accuracy using different A1 mask sizes in auditory attended runs, based on visual expectation. The left plot shows auditory unexpected trials, and the right plot auditory expected trials. * $p < 0.05$

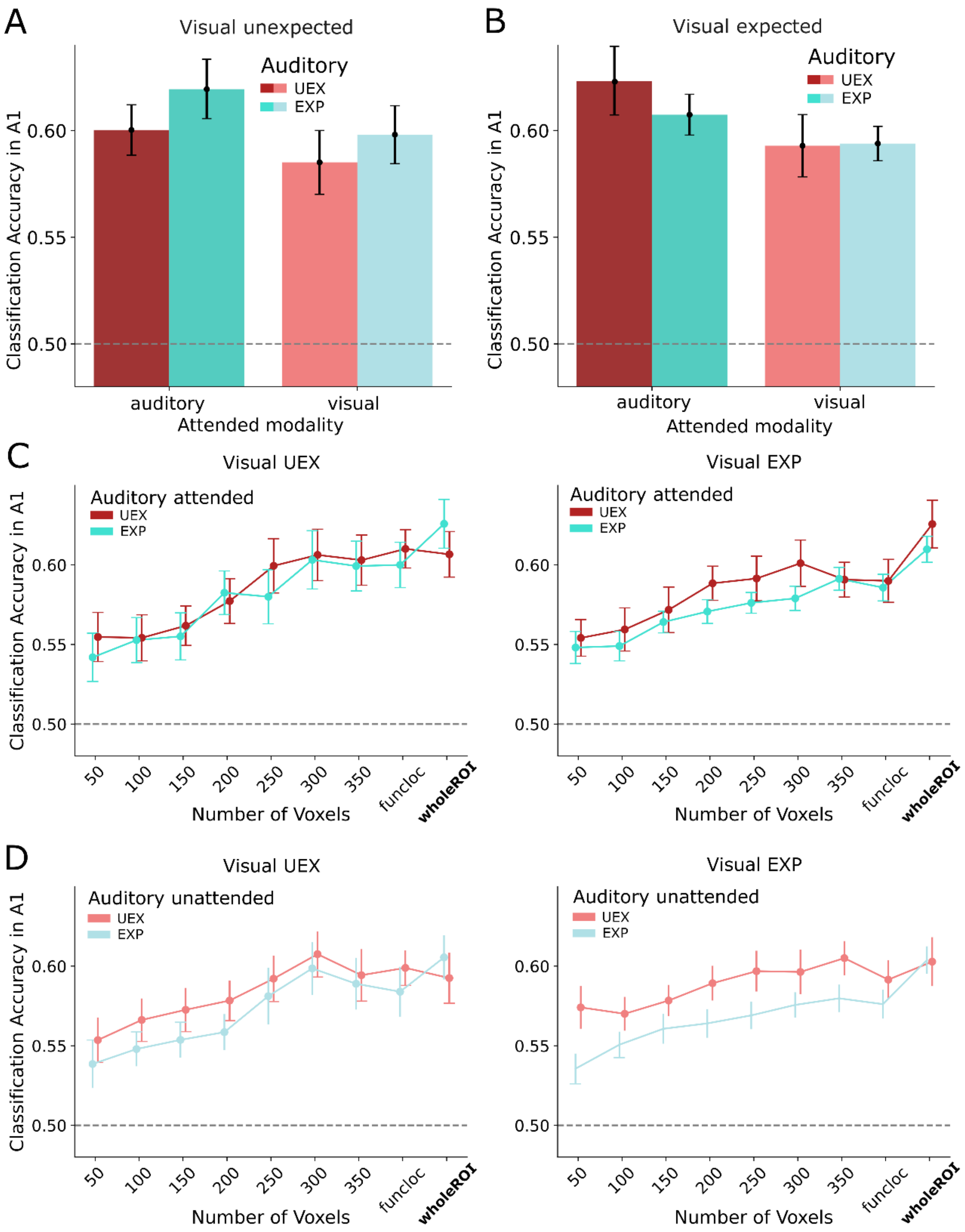


**Figure S3** (A) Classification accuracy in A1 based on attention and auditory expectation, in trials where the visual stimulus was unexpected. (B) Classification accuracy in A1 based on attention and auditory expectation, in trials where the visual stimulus was expected. (C) Classification accuracy using different A1 mask sizes in auditory attended runs, based on auditory expectation. The left plot shows visual unexpected trials, and the right plot visual expected trials. (D) Classification accuracy using different A1 mask sizes in visual attended runs, based on auditory expectation. The left plot shows visual unexpected trials, and the right plot visual expected trials.
