## Supplementary tables for "Attention and Explicit Knowledge Drive Predictive Sharpening in Early Visual Cortex"

***Table 1. Decoding Accuracy by ROI and mask size***

| ROI | ROI Size | n | Mean Accuracy | t-value | p-value | CI Lower | CI Upper | Significance |
|---|---|---|---|---|---|---|---|---|
| EVC | 50 | 24 | 0.524 | 2.65 | 0.014 | 0.505 | 0.543 | * |
| EVC | 100 | 24 | 0.538 | 4.36 | 0.000 | 0.520 | 0.556 | *** |
| EVC | 150 | 23 | 0.545 | 4.46 | 0.000 | 0.524 | 0.566 | *** |
| EVC | 200 | 23 | 0.555 | 4.95 | 0.000 | 0.532 | 0.579 | *** |
| EVC | 250 | 23 | 0.562 | 5.52 | 0.000 | 0.538 | 0.585 | *** |
| EVC | 300 | 23 | 0.568 | 5.94 | 0.000 | 0.544 | 0.592 | *** |
| EVC | 350 | 23 | 0.568 | 5.71 | 0.000 | 0.543 | 0.592 | *** |
| EVC | funcloc | 24 | 0.578 | 4.77 | 0.000 | 0.544 | 0.612 | *** |
| EVC | wholeROI | 25 | 0.562 | 4.15 | 0.000 | 0.531 | 0.593 | *** |
| A1 | 50 | 24 | 0.544 | 4.95 | 0.000 | 0.526 | 0.563 | *** |
| A1 | 100 | 22 | 0.555 | 5.05 | 0.000 | 0.533 | 0.578 | *** |
| A1 | 150 | 22 | 0.564 | 5.62 | 0.000 | 0.540 | 0.588 | *** |
| A1 | 200 | 21 | 0.577 | 6.99 | 0.000 | 0.554 | 0.601 | *** |
| A1 | 250 | 18 | 0.594 | 7.01 | 0.000 | 0.566 | 0.622 | *** |
| A1 | 300 | 18 | 0.604 | 7.79 | 0.000 | 0.576 | 0.633 | *** |
| A1 | 350 | 18 | 0.602 | 7.52 | 0.000 | 0.574 | 0.631 | *** |
| A1 | funcloc | 24 | 0.586 | 6.46 | 0.000 | 0.559 | 0.614 | *** |
| A1 | wholeROI | 25 | 0.603 | 8.31 | 0.000 | 0.577 | 0.628 | *** |

***Table 2. Decoding Accuracy Differences Between Attended and Unattended by ROI and Voxel Size***

| ROI | ROI Size | n | Mean Diff. | t-value | p-value | Cohen's d | CI Lower | CI Upper | Significance |
|---|---|---|---|---|---|---|---|---|---|
| EVC | 50 | 24 | 0.023 | 2.60 | 0.016 | 0.53 | 0.005 | 0.042 | * |
| EVC | 100 | 24 | 0.017 | 2.02 | 0.055 | 0.41 | 0.000 | 0.034 | |
| EVC | 150 | 23 | 0.029 | 3.59 | 0.002 | 0.75 | 0.012 | 0.045 | ** |
| EVC | 200 | 23 | 0.021 | 2.04 | 0.053 | 0.43 | 0.000 | 0.043 | |
| EVC | 250 | 23 | 0.028 | 2.33 | 0.029 | 0.49 | 0.003 | 0.053 | * |
| EVC | 300 | 23 | 0.033 | 2.66 | 0.014 | 0.56 | 0.007 | 0.059 | * |
| EVC | 350 | 23 | 0.041 | 3.36 | 0.003 | 0.70 | 0.016 | 0.066 | ** |
| EVC | funcloc | 24 | 0.028 | 2.21 | 0.037 | 0.45 | 0.002 | 0.054 | * |
| EVC | wholeROI | 25 | 0.025 | 1.93 | 0.066 | 0.39 | -0.002 | 0.053 | |
| A1 | 50 | 24 | -0.002 | -0.29 | 0.772 | -0.06 | -0.019 | 0.014 | |
| A1 | 100 | 22 | 0.007 | 0.84 | 0.409 | 0.18 | -0.010 | 0.025 | |
| A1 | 150 | 22 | 0.005 | 0.62 | 0.544 | 0.13 | -0.013 | 0.023 | |
| A1 | 200 | 21 | -0.005 | -0.55 | 0.591 | -0.12 | -0.024 | 0.014 | |
| A1 | 250 | 18 | 0.000 | 0.00 | 1.000 | 0.00 | -0.024 | 0.024 | |
| A1 | 300 | 18 | -0.001 | -0.07 | 0.948 | -0.02 | -0.028 | 0.027 | |
| A1 | 350 | 18 | -0.002 | -0.17 | 0.867 | -0.04 | -0.028 | 0.023 | |
| A1 | funcloc | 24 | -0.012 | -1.15 | 0.262 | -0.23 | -0.033 | 0.009 | |
| A1 | wholeROI | 25 | -0.020 | -2.21 | 0.037 | -0.44 | -0.039 | -0.001 | * |

***Table 3. Paired t-Tests for Visual Attended Decoding in EVC***

| ROI size | A predicted | t-value | df | p-value | $BF_{10}$ | Cohen's d | Significance |
|---|---|---|---|---|---|---|---|
| 50 | 0 | -0.37 | 23 | 0.698 | 0.457 | -0.10 | |
| 50 | 1 | 0.53 | 23 | 0.698 | 0.487 | 0.11 | |
| 100 | 0 | 1.56 | 23 | 0.934 | 0.801 | 0.38 | |
| 100 | 1 | -0.46 | 23 | 0.650 | 0.473 | -0.11 | |
| 150 | 0 | 0.46 | 22 | 0.675 | 0.482 | 0.09 | |
| 150 | 1 | -0.17 | 22 | 0.675 | 0.443 | -0.03 | |
| 200 | 0 | 0.59 | 22 | 0.720 | 0.513 | 0.13 | |
| 200 | 1 | -0.45 | 22 | 0.656 | 0.480 | -0.09 | |
| 250 | 0 | -0.08 | 22 | 0.468 | 0.439 | -0.02 | |
| 250 | 1 | -2.42 | 22 | 0.024 | 4.750 | -0.50 | * |
| 300 | 0 | -0.30 | 22 | 0.384 | 0.456 | -0.06 | |
| 300 | 1 | -1.58 | 22 | 0.129 | 1.286 | -0.32 | |
| 350 | 0 | -2.35 | 22 | 0.028 | 4.174 | -0.47 | * |
| 350 | 1 | -0.67 | 22 | 0.256 | 0.535 | -0.14 | |
| funcloc | 0 | 0.52 | 23 | 0.696 | 0.486 | 0.09 | |
| funcloc | 1 | -1.94 | 23 | 0.065 | 2.118 | -0.35 | |
| wholeROI | 0 | 0.30 | 24 | 0.616 | 0.439 | 0.05 | |
| wholeROI | 1 | -2.26 | 24 | 0.033 | 3.546 | -0.40 | * |

***Table 4. Paired t-Tests for Visual Unattended Decoding in EVC***

| ROI size | A predicted | t-value | df | p-value | $BF_{10}$ | Cohen's d | Significance |
|---|---|---|---|---|---|---|---|
| 50 | 0 | -0.61 | 23 | 0.726 | 0.508 | -0.18 | |
| 50 | 1 | -0.16 | 23 | 0.726 | 0.434 | -0.04 | |
| 100 | 0 | 0.38 | 23 | 0.353 | 0.459 | 0.09 | |
| 100 | 1 | 0.51 | 23 | 0.353 | 0.483 | 0.14 | |
| 150 | 0 | 0.26 | 22 | 0.649 | 0.452 | 0.06 | |
| 150 | 1 | -0.39 | 22 | 0.649 | 0.468 | -0.10 | |
| 200 | 0 | 0.59 | 22 | 0.370 | 0.512 | 0.14 | |
| 200 | 1 | 0.33 | 22 | 0.370 | 0.460 | 0.07 | |
| 250 | 0 | -0.99 | 22 | 0.833 | 0.676 | -0.20 | |
| 250 | 1 | 1.35 | 22 | 0.191 | 0.973 | 0.29 | |
| 300 | 0 | -1.58 | 22 | 0.936 | 0.774 | -0.30 | |
| 300 | 1 | 0.62 | 22 | 0.539 | 0.522 | 0.15 | |
| 350 | 0 | 0.16 | 22 | 0.877 | 0.442 | 0.04 | |
| 350 | 1 | -1.53 | 22 | 0.929 | 0.831 | -0.31 | |
| funcloc | 0 | -0.07 | 23 | 0.528 | 0.430 | -0.01 | |
| funcloc | 1 | 0.45 | 23 | 0.528 | 0.471 | 0.09 | |
| wholeROI | 0 | 0.46 | 24 | 0.326 | 0.464 | 0.11 | |
| wholeROI | 1 | 1.16 | 24 | 0.258 | 0.768 | 0.23 | |

***Table 5. Paired t-Tests for Auditory Attended Decoding in A1***

| ROI size | V predicted | t-value | df | p-value | $BF_{10}$ | Cohen's d | Significance |
|---|---|---|---|---|---|---|---|
| 50 | 0 | 0.48 | 23 | 0.703 | 0.476 | 0.11 | |
| 50 | 1 | 0.54 | 23 | 0.703 | 0.490 | 0.14 | |
| 100 | 0 | 0.07 | 21 | 0.745 | 0.447 | 0.02 | |
| 100 | 1 | 0.67 | 21 | 0.745 | 0.547 | 0.15 | |
| 150 | 0 | 0.30 | 21 | 0.696 | 0.465 | 0.07 | |
| 150 | 1 | 0.52 | 21 | 0.696 | 0.504 | 0.10 | |
| 200 | 0 | -0.22 | 20 | 0.831 | 0.465 | -0.05 | |
| 200 | 1 | 1.18 | 20 | 0.873 | 0.835 | 0.27 | |
| 250 | 0 | 0.65 | 17 | 0.806 | 0.586 | 0.18 | |
| 250 | 1 | 0.89 | 17 | 0.806 | 0.687 | 0.19 | |
| 300 | 0 | 0.10 | 17 | 0.876 | 0.488 | 0.03 | |
| 300 | 1 | 1.19 | 17 | 0.876 | 0.901 | 0.28 | |
| 350 | 0 | -0.04 | 17 | 0.554 | 0.486 | -0.01 | |
| 350 | 1 | 0.14 | 17 | 0.554 | 0.490 | 0.03 | |
| funcloc | 0 | 0.46 | 23 | 0.677 | 0.474 | 0.10 | |
| funcloc | 1 | 0.30 | 23 | 0.677 | 0.448 | 0.05 | |
| wholeROI | 0 | -1.03 | 24 | 0.314 | 0.679 | -0.19 | |
| wholeROI | 1 | 0.87 | 24 | 0.803 | 0.593 | 0.21 | |

***Table 6. Paired t-Tests for Auditory Unattended Decoding in A1***

| ROI size | V predicted | t-value | df | p-value | $BF_{10}$ | Cohen's d | Significance |
|---|---|---|---|---|---|---|---|
| 50 | 0 | 2.05 | 23 | 0.052 | 2.525 | 0.55 | |
| 50 | 1 | 0.71 | 23 | 0.243 | 0.539 | 0.20 | |
| 100 | 0 | 1.05 | 21 | 0.153 | 0.726 | 0.24 | |
| 100 | 1 | 1.27 | 21 | 0.153 | 0.910 | 0.32 | |
| 150 | 0 | 0.95 | 21 | 0.176 | 0.669 | 0.26 | |
| 150 | 1 | 1.18 | 21 | 0.176 | 0.822 | 0.28 | |
| 200 | 0 | 0.97 | 20 | 0.171 | 0.692 | 0.26 | |
| 200 | 1 | 1.53 | 20 | 0.142 | 1.245 | 0.47 | |
| 250 | 0 | 0.41 | 17 | 0.343 | 0.524 | 0.13 | |
| 250 | 1 | 1.54 | 17 | 0.142 | 1.321 | 0.49 | |
| 300 | 0 | 0.35 | 17 | 0.366 | 0.513 | 0.10 | |
| 300 | 1 | 1.10 | 17 | 0.287 | 0.821 | 0.34 | |
| 350 | 0 | 1.66 | 17 | 0.116 | 1.529 | 0.52 | |
| 350 | 1 | 0.19 | 17 | 0.427 | 0.494 | 0.06 | |
| funcloc | 0 | 0.65 | 23 | 0.262 | 0.519 | 0.17 | |
| funcloc | 1 | 0.82 | 23 | 0.262 | 0.580 | 0.21 | |
| wholeROI | 0 | -0.60 | 24 | 0.724 | 0.498 | -0.13 | |
| wholeROI | 1 | -0.07 | 24 | 0.724 | 0.423 | -0.02 | |
